## Supplementary material for "Inhibiting heme-piracy by pathogenic *Escherichia coli* using *de novo*-designed proteins": Table S4

**Extended Data Table 1: Cryo-EM data collection, refinement and validation statistics.** All datasets were collected with a zero-loss filtering slit width of 10 eV and with 70 frames per movie.

|  | ChuA-G7  (EMD-46916)  (PDB 9DIR) | ChuA-H3  (EMD-46917)  (PDB 9DIS) |
| --- | --- | --- |
| **Data collection and processing** |  |  |
| Magnification | 105kx | 105kx |
| Voltage (kV) | 300 | 300 |
| Electron exposure (e–/Å^2^) | 70.0 | 70.0 |
| Defocus range (μm) | -1.4 -0.5 | -1.4 -0.5 |
| Pixel size (Å) | 0.82 | 0.82 |
| Symmetry imposed | C1 | C1 |
| Initial particle images (no.) | 3,850,705 | 2,490,792 |
| Final particle images (no.) | 131,924 | 333,575 |
| Map resolution (Å)  FSC threshold | 2.97  0.143 | 2.51  0.143 |
| Map resolution range (Å) | 2.63 – 4.72 | 2.23 – 4.62 |
| **Refinement** |  |  |
| Initial model used (PDB code) | 3FHH | 3FHH |
| Model resolution (Å)  FSC threshold | 2.97  0.143 | 2.51  0.143 |
| Model resolution range (Å) | n/a | n/a |
| Map sharpening *B* factor (Å^2^) | 85.6 | 82.1 |
| Map-Model CC | 0.78 | 0.82 |
| Model composition  Non-hydrogen atoms  Protein residues  Ligands | 5870  755  0 | 5713  741  0 |
| Solvent | 0 | 0 |
| *B* factors (Å^2^)  Protein  Ligand | 108.92  n/a | 45.43  n/a |
| Solvent | n/a | n/a |
| R.m.s. deviations  Bond lengths (Å)  Bond angles (°) | 0.003  0.595 | 0.004  0.541 |
| Validation  MolProbity score  Clashscore  Poor rotamers (%) | 1.76  11.94  0.00 | 1.71  10.92  0.00 |
| Ramachandran plot  Favored (%)  Allowed (%)  Disallowed (%) | 97.05  2.95  0.00 | 97.41  2.86  0.00 |
